# snATAC-Express infers Gene Expression from Prioritized Chromatin Accessibility Peaks using Machine Learning

**DOI:** 10.1101/2025.07.25.666784

**Authors:** Margaret Brown, Alessandro Ferrari, Anne Dodd, Fang Shi, Vasantha L. Kolachala, Subra Kugathasan, Russell D. Wolfinger, Greg Gibson

**Author notes:** Email Addresses.

## Abstract

**Background:** Single cell multi-omic investigation opens-up new opportunities to understand mechanisms of gene regulation. Existing methods for inferring transcript abundance from chromatin accessibility fail to prioritize the most relevant peaks and tend to assume positive associations between ATAC peaks and RNA counts. We hypothesize that gene regulation can be modeled as a function of combined positive and negative interactions among peaks and that causal regulatory variants are enriched in the vicinity of the most critical peaks.

**Results:** A machine learning pipeline leveraging single nuclear multiomic transcriptome and chromatin accessibility data is developed to model gene expression as a function of ATAC peak intensity. Multiome data was available for 18 immune cell types from 29 donors, 19 with Crohn’s disease. The pipeline aggregates results from three machine learning approaches (random forest regression, XGBoost, and Light GBM) as well as linear regression to identify which ATAC peaks contribute to explaining variation among donors and cell types in pseudobulk gene expression. The coefficient of determination with cross-validation was used to identify robust models which typically explain between 5% and 40% of transcript abundance, utilizing on average 47% of the ATAC peaks, representing a significant gain in predictive accuracy. The most important peaks are enriched in GWAS variants for inflammatory bowel disease and the autoimmune disease systemic lupus erythematosus, but not for rheumatoid arthritis.

**Conclusion:** Atlanta Plots visualize the proportion of ATAC peaks contributing to a predictive model of gene expression as well as the proportion of variance explained by the model. Software implementing our pipeline, “snATAC-Express”, is freely available on GitHub.

## Background

Genome wide association studies (GWAS) have identified thousands of loci associated with heritable traits [1,2]. It is thought that the vast majority of variants influence gene expression since lead polymorphisms are located outside exons and have enriched overlap with eQTL [3]. In the post-GWAS era, comparative and descriptive omic analyses have been performed to obtain better insight into the properties of likely causal variants, and recently, single cell omics studies have further increased granularity [3, 4]. In addition to single cell RNA-sequencing (scRNA-seq), which provides transcriptomic data, snATAC-seq (single nucleus assay for transposase accessible chromatin sequencing) is now able to identify open chromatin that marks plausible regulatory elements such as enhancers, also in nuclei isolated from single cells [5–7]. The notion that many GWAS causal variants lie within open chromatin peaks and influence transcription of nearby genes [8,9] gives rise to two testable hypotheses: (i) that some differentially accessible chromatin peaks are more important than others for gene regulation; and (ii) that those peaks will preferentially lie in close proximity to GWAS lead variants. Here we test these hypotheses in the context of three immune disorders using a novel machine learning-driven strategy for modeling gene expression as a function of joint contributions of accessible chromatin peaks.

Joint single nuclear RNA-seq and ATAC-seq analyses on the same cells, hereafter termed multiomics, is beginning to be deployed to characterize and better understand gene regulatory networks [10]. Various tools and databases now support annotation of potential transcription factor binding sites (TFBS) within genomic coordinates of ATAC-seq peaks, which may in turn lie in the vicinity of transcribed DNA. If the TFBS are also coincident with the credible set of a GWAS peak, and the nearby gene is differentially expressed, multi-omic data has the potential to provide functional annotations to GWAS tagged loci [11–13]. While these methods link peaks in ATAC-seq data with genes by virtue of proximity, functional inference is further supported if there is evidence that an ATAC peak is relevant to a gene’s expression. Many studies assume that increased chromatin accessibility directly correlates with increased gene expression. However, substantial evidence indicates that chromatin accessibility can also be linked to reduced gene expression due to mechanisms like active repression [14–16]. Regression methods that allow for joint positive and negative influences as well as interactions among peaks thus hold promise for more accurate prediction of transcript abundance from ATAC-seq data.

Two of the currently most popular analytical pipelines, ArchR and Signac, both implement a framework for inference of gene expression which computes a “Gene Score” or a “Gene Activity Score” by summing the chromatin accessibility within a nearby a gene [17, 18]. Cicero, another software application which computes a “Gene Activity Score”, focuses on peaks which are co-accessible instead of considering the total chromatin accessibility [19], but also assumes that increased chromatin accessibility indicates increased gene expression.

Few studies have assessed to what extent individual ATAC-seq peaks explain gene expression, and nor have many studies systematically prioritized which ATAC peaks may be more or less important for explaining gene expression. This is a particularly striking omission given that the rationale for mapping differentially accessible regions (DAR) is that they are thought to be at least partially responsible for differential expression. Thus, a model expressing transcript abundance as a function of a weighted sum of accessibility at only the fraction of chromatin peaks that actually mediate gene expression ought to be superior to a model based either on total open chromatin or just co-accessible peaks. It follows logically that only those peaks that do explain some fraction of transcript abundance are likely to harbor polymorphisms associated with gene expression (and visible traits), and hence that such polymorphisms ought to be enriched in the functionally relevant peaks, rather than simply in the vicinity of the transcription unit.

Recently, two studies have filled this gap in the literature by utilizing joint single nucleus RNA and ATAC-seq to predict gene expression from chromatin accessibility. Sakaue et al. (2024) developed SCENT, a nonparametric statistical model based on Poisson regression for this purpose, also identifying expression-associated enhancers enriched in fine-mapped causal variants [20]. Mitra et al. (2024) similarly proposed a Poisson regression method named SCARlink [21]. Both SCENT and SCARlink demonstrated improved linking of chromatin accessibility with gene expression and observed an enrichment of trait-associated variants in prioritized regions of chromatin accessibility. While these methods demonstrate examples of uncovering unique biology between trait associated variants and important chromatin accessible peaks, neither method performed feature selection on peaks included in their models, nor determined how much their prioritized chromatin accessible regions explained gene expression compared to non-prioritized chromatin accessible regions.

To address this need, we present “snATAC-Express”, a pipeline which trains machine learning models on snATAC-seq data to infer gene expression measured by snRNA-seq and to prioritize expression-relevant peaks. The dataset comprises single nuclear multiomic data of circulating immune cells from a small cohort of Crohn’s disease and healthy individuals. We observe that machine learning models outperform linear regression models, confirming that the relationship between chromatin accessibility and gene expression is more complex than simple correlation between increased accessibility and increased expression. Ensembling the models, we quantify how much chromatin accessibility can explain gene expression and observe that the abundance of certain transcripts can be explained by chromatin accessibility more than others, somewhat independent of the number or proportion of peaks in the model. For each gene, we prioritize which peaks are better predictors of gene expression than others and utilizing a rigorous cross-validation approach demonstrate that only a subset of ATAC peaks are relevant to gene expression rather than considering all peaks within a cis-regulatory region. We determine the contribution each peak makes to gene expression prediction and determine that a substantial number of ATAC peaks with increased accessibility may be associated with reduced gene expression. We evaluate the landscape of these peaks in the context of co-accessibility and by annotating tagged GWAS variants. Our analysis confirms an enrichment of autoimmune GWAS variants in ATAC peaks that are identified as important for predicting immune gene expression.

## Results

### Predicting gene expression from ATAC-seq data

To evaluate the predictive capacity of ATAC-seq data for gene expression, we trained a series of models on pseudobulked snATAC-seq data and assessed their performance using k-fold cross validation R^2^ (see Methods). For each gene of interest, we individually modeled gene expression, which was measured as the pseudobulk log base 2 normalized expression of the gene in a sample and cell type. The ATAC peaks considered for each gene were identified by finding peaks that overlapped the gene’s cis-regulatory region, heuristically defined as 100kb upstream and downstream of the transcription start site (TSS) and the transcription end site (TES) respectively and including all introns and exons (see Methods) [22]. Our methodology is based on the idea that ATAC peaks are mechanistically involved with gene expression, and that chromatin accessibility is associated with gene expression [23]. There are two sources of biological variance contributing to both chromatin accessibility and transcript abundance: among individual differences, and cell-type specific differences. With 29 donors and 18 peripheral blood cell types, there are 342 vectors of data per gene model.

The modeling strategy is illustrated in Figure 1A, where six peaks within the cis-regulatory region of a gene hypothetically exhibit varying intensities and may collectively explain variable levels of expression of a nearby gene. The first peak is unlikely to have a functional role since Peak 1 is not correlated with gene expression: increased chromatin accessibility is observed at the extreme levels of no gene expression or high gene expression. By contrast, Peak 2 exhibits a negative correlation with gene expression, having chromatin accessibility in samples with lower gene expression. Peaks 3 and 4 are co-accessible, showing similar levels of accessibility and positive correlation with gene expression. Peak 5 is also positively correlated with gene expression, and in this case, varying levels of chromatin accessibility correlate with the varying levels of gene expression. Peak 6 is co-accessible with Peak 5 and is also positively correlated with gene expression, however this peak is relatively less accessible than Peak 5. In reality, many more peaks are observed near most genes, there may be many more possible relationships, and variable proportions of the peaks may correlate with gene expression. Due to this complexity we have opted to use machine learning models to capture these nuanced relationships effectively, given their ability to capture complex, nonlinear relationships [24]. We selected a total of 502 genes tagged from GWAS for three traits: inflammatory bowel disease (IBD), rheumatoid arthritis (RA), and systemic lupus erythematosus (SLE) (see Methods, Figure 1B). These traits were chosen due to the high likelihood that pathology traces in part to immune cells [25–28].

**Figure 1.**
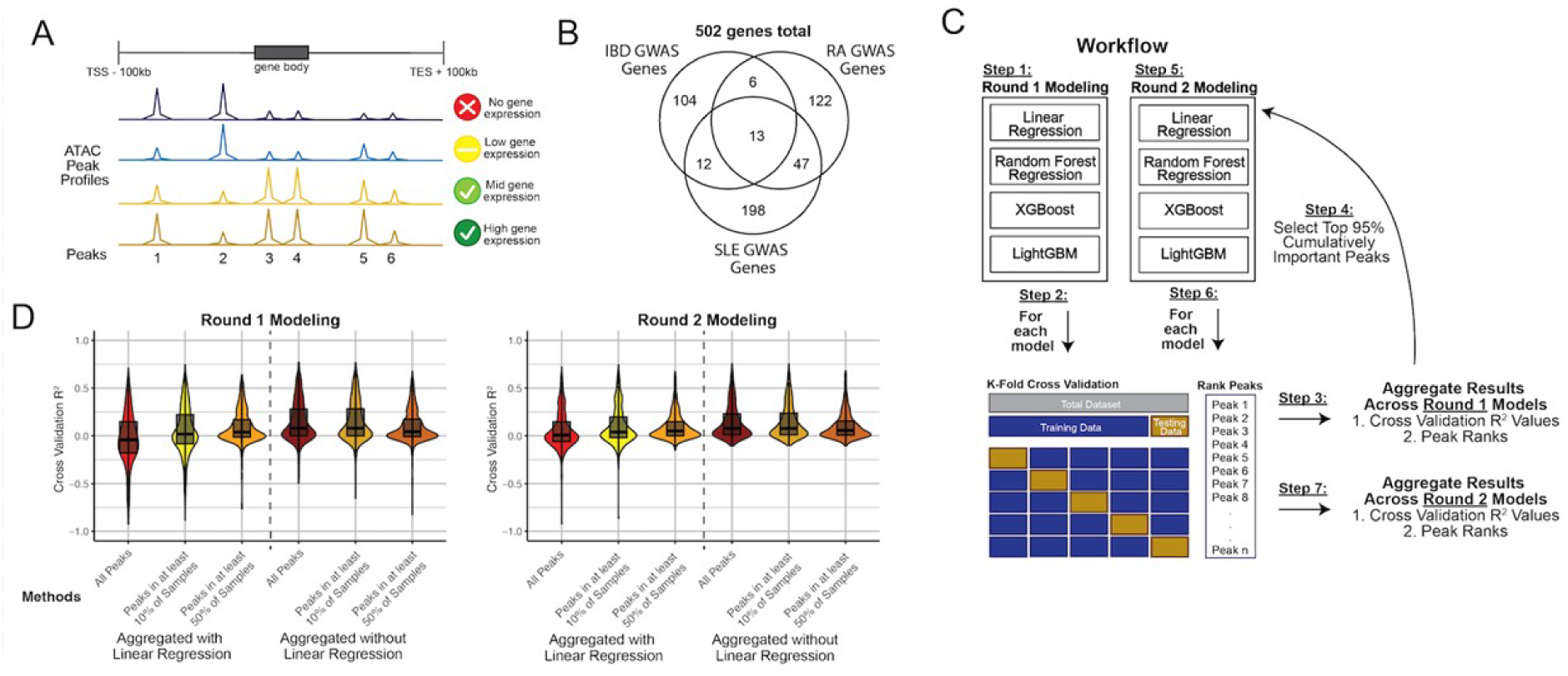
Model overview. (A) Depiction of how chromatin accessible peaks within a gene’s cis-regulatory region may or may not correlated clearly with gene expression. (B) Venn diagram of genes tagged by variant associations from GWAS for three traits: IBD, RA, and SLE. (C) Overview of predictive modeling pipeline. Round 1 models are performed for four prediction strategies linear regression, random forest regression, XGBoost, and LightGBM, which each undergo k-fold cross validation and peak rankings within each fold. The top 95% cumulatively important peaks are identified and used for training for a second round of predictive modeling. (D) Ensembled results of each gene for Round 1 and Round 2 models, aggregated with and without linear regression, using all peaks mapped to the gene, peaks in at least 10% of samples which mapped to the gene, and peaks in at least 50% of samples mapped to the gene. IBD, Inflammatory Bowel Disease; RA, Rheumatoid Arthritis; SLE, Systemic Lupus Erythematosus.

We utilized four distinct models: linear regression, random forest regression (RFR), XGBoost (XGB), and LightGBM (LGBM) (see Methods). For each gene, we employed two rounds of modeling, both followed by k-fold cross validation and peak ranking (Figure 1C). The initial round of modeling is implemented to determine the importance of each ATAC peak. The top 95% cumulatively important peaks were identified from the aggregated peak ranks across the models, and then used for a second round of modeling. Furthermore, these modeling strategies were implemented three times with different numbers of peaks according to how many of the samples they are observed in. The first implementation included all peaks in the cis-regulatory region for the first round of modeling. The second implementation was slightly more restrictive as it only included peaks present in at least 10% of the pseudobulked samples, while the third implementation included peaks present in at least 50% of pseudobulked samples and hence which are more broadly active. While each machine learning model performed comparably, linear regression consistently underperformed across all scenarios (Supplementary Figure 1). Consequently, the results were evaluated by aggregating k-fold cross validation R^2^ values (and peak ranks) with and without linear regression, shown in Figure 1D. For both the first and second round modeling outcomes, using all peaks within the region outperformed models trained on peaks present in at least 50% of pseudobulked samples (Welch Two Sample t-test, p-value = 6.0e-5 and 7.1e-5 for the first and second rounds, respectively; see Table 1). Similarly, models trained on peaks in at least 10% of pseudobulked samples within the region outperformed models trained on peaks in at least 50% in pseudobulked samples (Welch Two Sample t-test, p-value = 1.2e-4 and 6.3e-5 for the first and second rounds, respectively). No significant difference was observed between training on all peaks and training on peaks present in at least 10% of pseudobulked samples. Additionally, there was no discernible difference between the first and second rounds of modeling, alleviating concerns that the second round might overfit due to biases from the first round. Collectively these results indicate that modeling with a subset of prioritized ATAC peaks can explain gene expression as effectively as using all ATAC peaks. The corollary is that all peaks within a cis-regulatory region of a gene may not necessarily contribute to gene expression. As a result, we focused on modeling results from the second round of modeling trained on peaks in at least 10% of samples. These results are from the machine learning models which were aggregated, while excluding linear regression.

**Table 1.** Aggregated R^2^ Values Computed as 1-(RSS/TSS) from K-Fold Cross Validation.

| Linear Regression Inclusions | Peak Filters | Round 1 Modeling Aggregated K-Fold Cross Validation $R^2$ Values | | Round 2 Modeling Aggregated K-Fold Cross Validation $R^2$ Values | |
| --- | --- | --- | --- | --- | --- |
|  |  | Mean | Median | Mean | Median |
| Include Linear Regression in Aggregation | All Peaks | 0.00 | -0.06 | 0.00 | 0.00 |
| | Peaks in $\geq 10\%$ of Samples | 0.03 | 0.01 | 0.01 | 0.04 |
| | Peaks in $\geq 50\%$ of Samples | 0.09 | 0.04 | 0.01 | 0.05 |
| Exclude Linear Regression in Aggregation | All Peaks | 0.15 | 0.08 | 0.14 | 0.08 |
| | Peaks in $\geq 10\%$ of Samples | 0.15 | 0.08 | 0.14 | 0.08 |
| | Peaks in $\geq 50\%$ of Samples | 0.10 | 0.04 | 0.10 | 0.06 |

### Machine learning models perform better than ArchR

To evaluate how our models perform against comparable strategies, we assessed our model’s ability to explain gene expression variance as well as how well the predicted gene expression values are correlated with real values in comparison to the “gene score” method in ArchR [17]. ArchR’s “gene score” considers three major components: 1) accessibility within the entire gene body; 2) an exponential weighting function accounting for activity of putatively distal regulatory elements; and 3) imposing gene boundaries to minimize usage of unrelated regulatory elements. Notably, it does not attempt to model contributions of individual chromatin accessibility peaks.

We compared our models to ArchR by computing both the squared correlation coefficient *r*^2^ value and the cross-correlation coefficient of determination R^2^ value. First, we evaluated the linear Pearson correlation between the predicted and actual gene expression; the square of this value (ranging from 0 to 1) is often taken as a measure of the proportion of variance explained by the model. However, a more robust measure of goodness of fit that takes into account differences unrelated to the ATAC peaks between the prediction and validation datasets is to compute the coefficient of determination, 1-(RSS/TSS), where RSS is the residual sum of squares and TSS is the total sum of squares, resulting in an R^2^ value that can range from -1 to 1 (see Methods). This is the R^2^ value calculated to evaluate the cross validation of machine learning models in Figure 1 and Table 1, and assesses the variance explained by the model versus the total variance in the data as an indicator for how well the model explains the variance in the data.

ArchR’s gene scores were compared against the real gene expression values, depicted in Figure 2. Results for the four individual models in both rounds of fitting are shown in Supplementary Figure 1, whereas Figure 2 reports model performance after aggregating the results (see Methods). Although visual inspection of violin plots in Figure 2A implies that the squared correlation coefficient distributions were similar across all implementations, the machine learning models aggregated without linear regression trained on all peaks and peaks in at least 10% of samples consistently outperformed ArchR’s gene score values. The linear least squares *r*^2^ value of machine learning models for all peaks had mean and median values of 0.16 and 0.10 respectively, models trained on peaks in at least 10% of samples had mean and median values of 0.17 and 0.10, while ArchR had mean and median values of 0.15 and 0.08, respectively (Supplementary Table 1). Based on these results, the machine learning models modestly outperformed ArchR’s gene scores.

**Figure 2.**
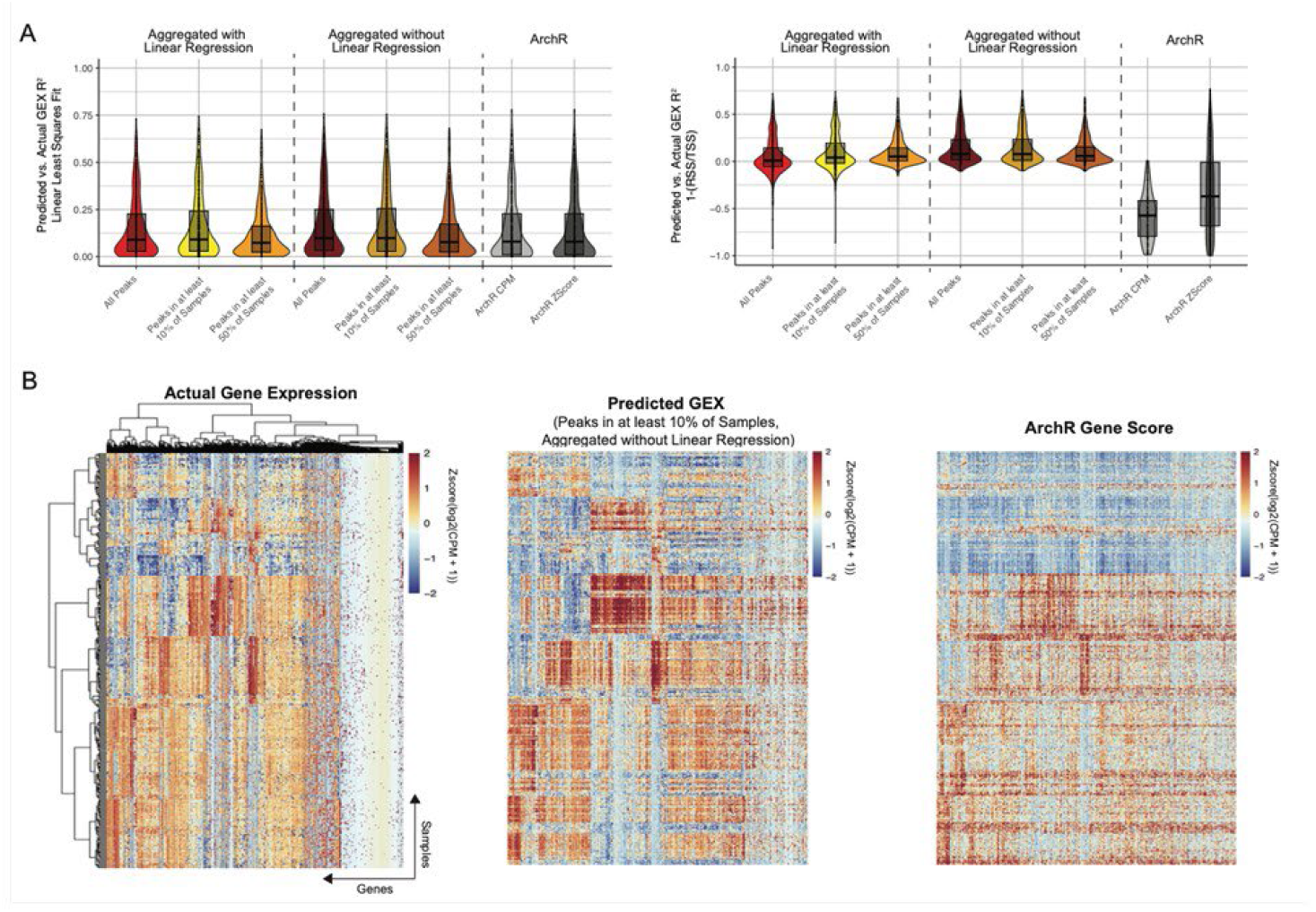
Machine learning predictions compared with ArchR’s Gene Scores. (A) Comparison of predicted gene expression values versus true gene expression values across all models, measured by linear least squares R2 value (left). Comparison of predicted gene expression values versus true gene expression values across all models, measured by R2 value computed as 1-(RSS/TSS) (right). (B) Heatmaps illustrating the actual gene expression (left), predicted gene expression from Round 2 modeling of the top 95% cumulatively important peaks present in at least 10% of samples (middle), and ArchR’s predicted gene scores (right). Values are z-scored log2(CPM+1). RSS, residual sum of squares; TSS, total sum of squares; GEX, gene expression; CPM, counts per million.

Turning to the cross-validation coefficient of determination R^2^ values, machine learning models outperformed ArchR’s gene scores more meaningfully. Both versions of models trained on all peaks plus peaks in 10% of samples having mean and median values of 0.14 and 0.08 respectively, whereas ArchR’s mean and median values fell below 0 with mostly negative R^2^ values. This indicates that in validation sets, ArchR’s gene score values perform worse than the average of the observed training data.

This improvement in performance is not necessarily surprising since gene expression is not included in the modeling performed by ArchR, but it nevertheless highlights the gains to be made by explicit multivariable modeling. In Figure 2B, the true gene expression values, compared to the predicted gene expression values from the best performing machine learning model pipeline trained on peaks in at least 10% of samples, plus ArchR’s gene score values are depicted in heatmaps across each sample. The actual gene expression values are clustered by both sample and gene, and the same order of gene and samples is retained for the heatmaps for both prediction methods. Improved model fit is particularly notable in the top right-hand quarters of the heatmaps. Overall, the machine learning model predictions more closely reflect the true gene expression values than ArchR’s gene scores.

### A subset of ATAC-seq peaks explain gene expression

After assessing various model implementations, we recognized that using a subset of features within a gene’s cis-regulatory region explained as much gene expression variance as using all peaks within the region, sometimes more. To explore this further, we assessed the number of peaks retained per gene from the best performing implementation, namely filtering for peaks in at least 10% of samples, ensembling only the machine learning models, after identifying the top 95% cumulatively important peaks after averaging the peak importance ranks (see Methods). For the GWAS associated genes, we identified enriched pathways within this gene set and assessed the proportion of peaks within each gene which was used to predict gene expression, as well as the gene’s k-fold cross validation R^2^ value (computed as 1-RSS/TSS).

Figure 3 introduces “Atlanta Plots” for visualization of these results. The gray “buildings” represent the total number of peaks within the gene’s cis-regulatory region, the green “trees” represent the actual number of peaks included in the model, and the blue “lake” represents the amount of gene expression explained. The individual plots corresponding to “downtown” and “midtown” gene neighborhoods are clustered according to enriched pathway, with the order of genes in each case from left to right reflecting the average variance explained per peak included in the model.

**Figure 3.**
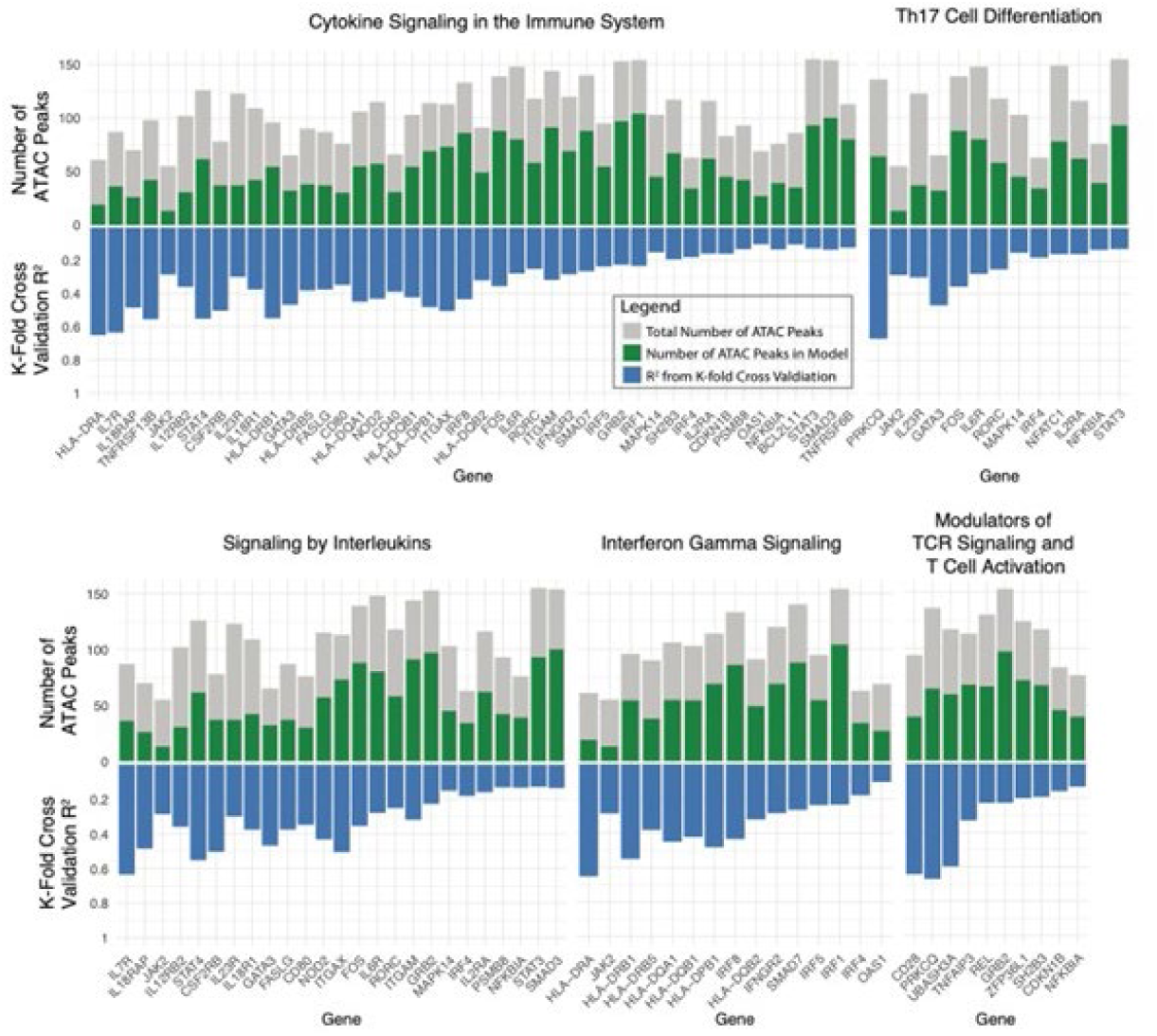
Atlanta plots show that a subset of peaks explain gene expression. Atlanta plots of GWAS tagged genes from IBD, SLE and RA which have been grouped by pathway analysis. The gray bars represent the total number of peaks within the gene’s cis-regulatory region, and the green bars represent the number of peaks in the region used to trained on for gene expression prediction. The blue bars represent the amount of gene expression explained, as the ensembled R^2^ value from k-fold cross validation. Genes are clustered by pathways and arranged from left to right according to the ratio of the proportion of peaks included to the amount of variance explained.

We assessed pathways associated with immune function and inflammation, including cytokine signaling, Th17 cell differentiation, signaling by interleukins, interferon gamma signaling, and modulators of T cell receptor (TCR) signaling and T cell activation [29–33]. We also assessed all genes identified by GWAS for each of the three diseases, IBD, SLE and RA, shown in Supplementary Figure 2A-C. While there is not a clear pattern as to whether inclusion of a greater or lesser proportions of peaks is better at explaining gene expression (Supplementary Figure 3), these results show that there is considerable variance among genes in the degree to which the expression is able to be explained by ATAC peaks. A notable counterintuitive trend is for the number of accessible chromatin peaks to increase as the coefficient of determination decreases, whereas there is no relationship between the proportion of peaks retained and variance explained (Supplementary Figure 3). These results show that some genes are inherently noisier, have lower heritability, or may be more susceptible to additional epigenetic or other mechanisms [34]. In addition, they confirm the intuition that not all peaks within a gene’s cis-regulatory region are relevant to that gene’s expression. In fact, for all genes we assessed with an R^2^ value > 0, an average of 47% of peaks within a gene’s cis-regulatory region were able to explain the same amount (if not more) gene expression than using all peaks within the region.

One gene of interest, *HLA-DRA*, was observed to be a gene tagged by GWAS for all three immune diseases and plays a role in both cytokine signaling and interferon gamma signaling [35, 36]. Training on 19 of 61 of its peaks within its cis-regulatory region was able to explain approximately 65% of the gene expression. In Supplementary Figure 4, the ATAC profile for *HLA-DRA* is shown, with the peaks used in the model highlighted in green underneath the track plot which included peaks upstream, within the gene body, and downstream, even overlapping some neighboring genes including *BTNL2* and *HLA-DRB5* and excluding peaks closer to *HLA-DRA’s* gene body. This example demonstrates that proximity is not a reliable indicator for identifying which peaks are relevant to gene expression and suggests that utilizing all identified peaks within the region likely includes noisy peaks which may not be an accurate reflection of the biology.

### Chromatin accessibility does not always positively correlate with increased gene expression

To determine how many ATAC peaks were correlated with gene expression in discordant directions, we used Shapley values from our best performing ensembled model trained on peaks present in at least 10% of samples to determine how much each peak contributed to gene expression prediction, and compared these Shapley values against the ATAC peak values to identify whether they are positively or negatively correlated (see Methods) [37–39]. In addition, we assessed each ATAC peak individually and compared accessibility values to its associated gene’s expression, to determine whether the peak is positively or negatively correlated with the expression. Because Shapley values consider the peaks in the context of the model, they can account for nonlinear relationships, interaction effects and redundancy, thus indicating a potentially more accurate estimation of how each peak contributes to gene expression prediction. For the 500 genes we modeled, we evaluated 29,565 gene-peak pairs, with an average of 59 peaks trained per gene.

Using the Shapley values, we found that approximately 50% of peaks contributed to a reduction of predicted gene expression values (Table 2), suggesting that about half of the peaks used in the model are negatively associated with gene expression. By contrast, evaluating the gene expression against each individual ATAC peak trained in the models, only 15% of peaks were negatively correlated with gene expression, indicating that marginal linear correlations of each peak’s accessibility versus gene expression considered in isolation overestimates the correlation between increased accessibility and increased gene expression. Supplementary Table 2 extends this analysis by trait (IBD, SLE, RA), showing consistent patterns across disease contexts.

**Table 2.**
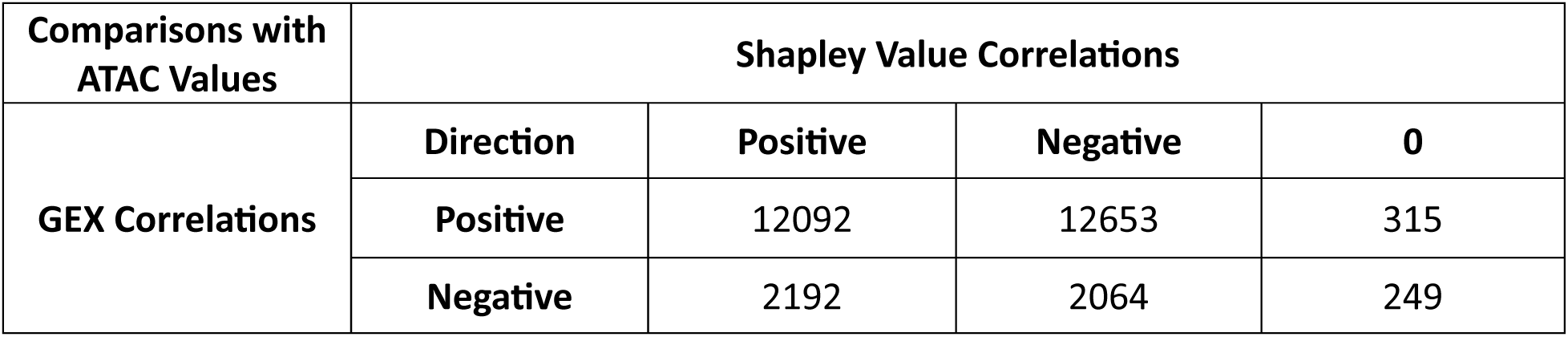
Direction of ATAC Peaks Impact on GEX Prediction.

Although we identified 29,565 gene-to-peak pairs, only 20,712 unique peaks were included, indicating that a number of ATAC peaks were trained for more than one gene. Indeed, 27% (5,533) of peaks were included in gene expression prediction models for more than one gene, consistent with at least one quarter of chromatin accessible regions moderating the expression of more than one gene at a time (Supplementary Table 3). To quantify the regulatory complexity of these shared peaks, we documented the directionality and magnitude of their Shapley values across different gene contexts. Of the 5,533 peaks paired with multiple genes, 2,404 (43%) peaks acted consistently in the same direction for all associated genes according to their Shapley values: 1,178 (21.3%) showing exclusively positive contributions with median Shapley values of +0.42 (IQR 0.18 to 0.71), and 1,226 (22.2%) showing exclusively negative contributions with median values of -0.38 (IQR: -0.65 to -0.15). Strikingly, 3,129 peaks (57%) exhibited context-dependent regulatory behavior, positively contributing to at least one gene’s predicted expression while negatively contributing to others. These bi-directional peaks showed a median absolute Shapley value difference of 0.84 between their positive and negative contributions (IQR: 0.52 to 1.31), indicating substantial regulatory switching rather than minor fluctuations around zero. Such bidirectional regulatory capacity is consistent with known mechanisms including: (i) competitive binding between activators and repressors at shared motifs, (ii) promoter-specific interpretation of enhancer signals through distinct cofactor recruitment, and (iii) three-dimensional chromatin interactions that bring the same regulatory element into different gene-specific regulatory hubs. These findings underscore why simple models assuming uniform positive correlations between accessibility and expression fail to capture the true complexity of gene regulation.

One gene of interest, *CD40*, was observed to harbor GWAS tagged variants within trained peaks for all three of our traits of interest: IBD, SLE and RA. This gene is primarily expressed in B cells (Figure 4A), with potentially increased expression in Activated B Cells of Crohn’s disease donors compared to Healthy donor Activated B Cells. A total of 66 ATAC peaks in our multiome dataset mapped to the cis-regulatory region of *CD40*, and after feature selection, 30 peaks were retained for training which explain approximately 40% of *CD40* expression. Upon assessing Shapley values for these peaks and their contributions to *CD40* predicted expression, the second, third and fourth largest contributing peaks were observed to be peaks harboring GWAS variants for IBD, SLE, and RA (Figure 4B). The second peak, chr20:46118998-46119498 harbors SNP rs4810485, a tagged GWAS variant associated with both SLE and RA [40, 41]. The third peak, chr20:46120489-46120989 harbors SNP rs4239702, a tagged GWAS variant associated with RA [42]. Finally, the fourth largest contributing peak, chr20:46111121-46111621 harbors SNP rs6074022, a tagged GWAS variant associated with IBD [43]. The SNPs are at different allele frequencies but in high LD (D’ > 0.95) according to LDLink [45]. Each of these SNPs have also been previously reported to be single cell eQTL in circulating immune cells on scQTLbase [44]. We assessed whether these peaks are likely to contribute to increased or decreased prediction values for gene expression and observed positive correlations between their chromatin accessibility and increased gene expression (Figure 4C). Upon closer inspection of the location of the variants, we show the ATAC track plots for *CD40* and mark the peaks containing the variants as red lines (Figure 4D). One variant is present in an ATAC peak upstream of *CD40*, while the other two are intragenic. This strategy of prioritizing which peaks contribute to gene expression and annotating them with GWAS variants is suggestive of function, but experimental validation provides stronger evidence and the ability to test computationally generated hypotheses.

**Figure 4.**
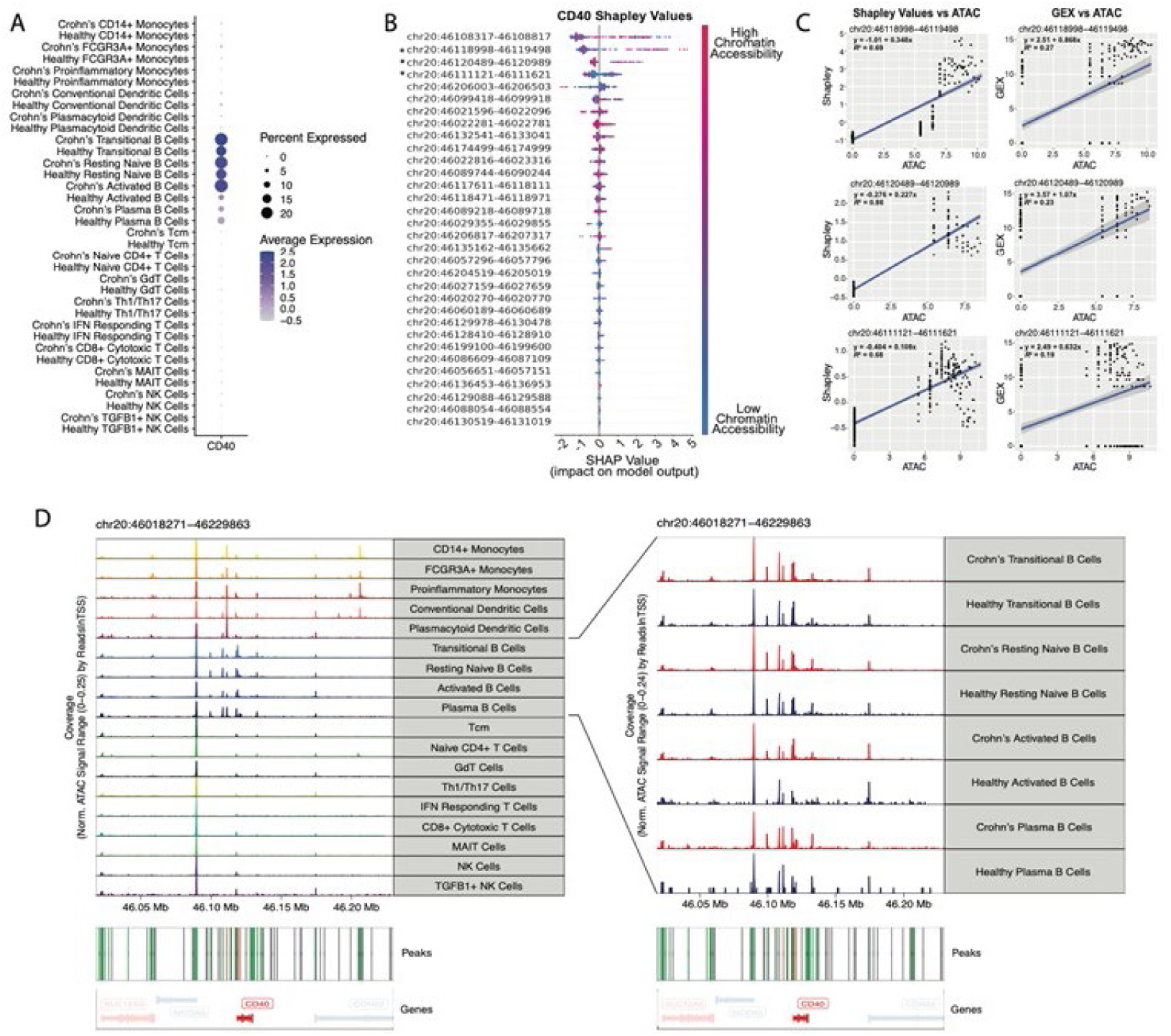
Assessment of Shapley values for ATAC peak contributions for CD40. (A) Dotplot showing the gene expression for CD40, primarily in B cell populations. (B) Beeswarm plot depicting the Shapley values for each ATAC peak per sample, ordered by the absolute value of the mean Shapley values per peak. Peaks marked with an asterisk harbor a GWAS variant associated with either SLE, IBD, and/or RA. (C) Scatter plot showing the correlation between the accessibility values of three ATAC peaks harboring GWAS variants for CD40 versus their Shapley values (left) and gene expression (right). ATAC accessibility values and gene expression values are pseudobulked, log2(CPM+1). (D) Track plots for CD40, with peaks not used for training shown in grey lines, and peaks which are used for training in the red and green lines. Red lines indicate peaks which harbor a GWAS variant.

### Important ATAC-seq peaks tend to be co-accessible

To assess the co-accessibility of peaks identified to be important for explaining gene expression, a Spearman correlation was computed between each peak pairwise with all other peaks trained for predicting a gene’s expression. For each gene, the top 95% cumulatively important peaks which were present in at least 10% of samples was evaluated and were divided into three “tiers”: the top third cumulatively important peaks in Tier 1, the middle third important peaks in Tier 2, and the least third important peaks as Tier 3 (see Methods). Across 500 genes influencing all three diseases, the vast majority (398) had moderate co-accessibility (average Spearman correlation between 0.34 and 0.67) among the Tier 1 peaks, whereas co-accessibility was on average lower in the Tier 2 and Tier 3 genes for more than half of these. Only 67 (13%) of the genes had low co-accessibility across all three Tiers. See Supplementary Table 4 for a more in-depth characterization of these patterns.

Three examples are depicted in Figure 5. Panel A shows the co-accessibility heatmap for 19 peaks used for training gene expression prediction for the *HLA-DRA* gene. The first 6 peaks in Tier 1 have an average Spearman correlation value of 0.43, while the peaks in Tiers 2 and 3 have correlations of 0.18 and 0.19 respectively. The other two examples are *GALC* which has similar levels of low-to-moderate co-accessibility across the entire locus, and *BANK1* for which there appears to be a gradient with low co-accessibility only observed in the third tier. Few genes were observed to have any tiers with a high average co-accessibility (Spearman correlation of > 0.67, 13 genes in total). These findings indicate that the peaks which are important for explaining gene expression tend to be co-accessible, but also show that many important peaks are independent of the others that contribute to gene regulation.

**Figure 5.**
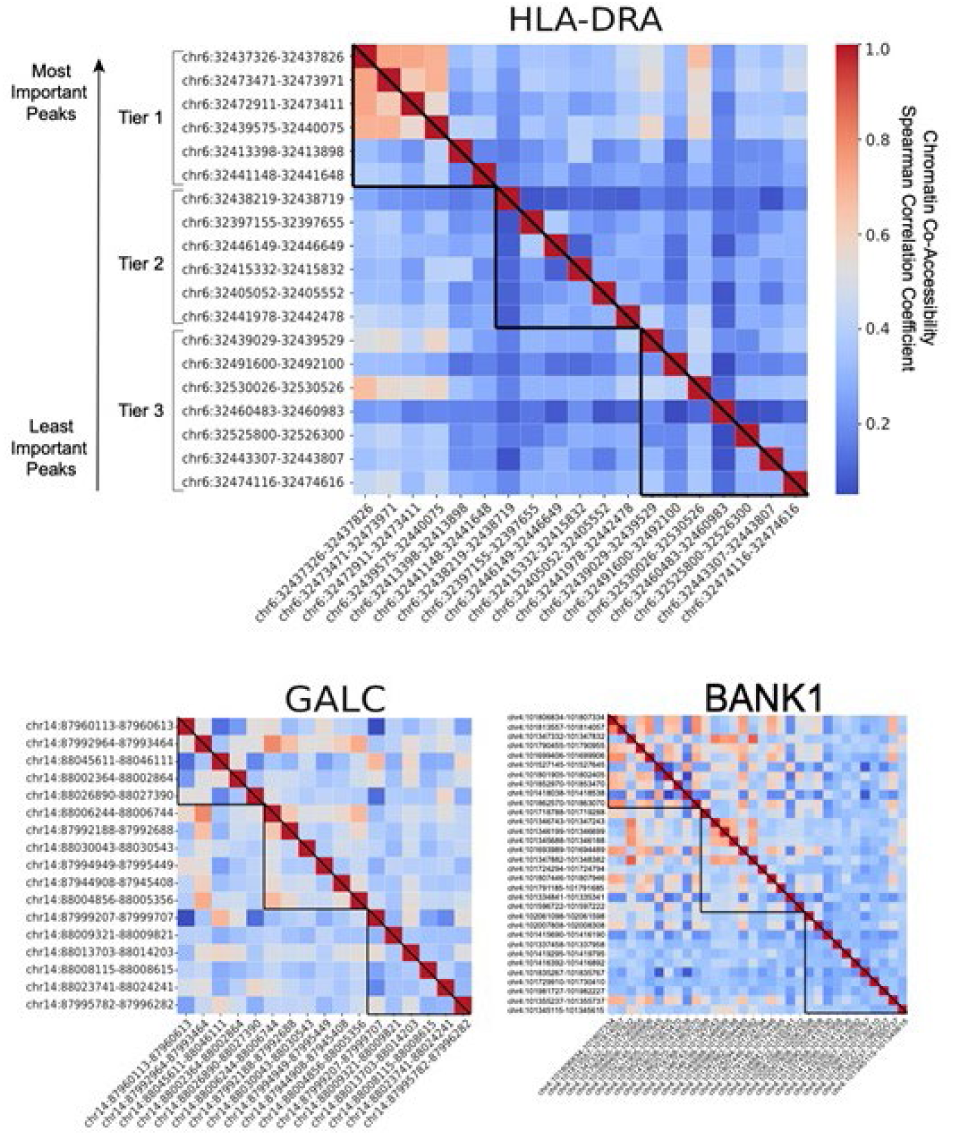
Co-accessibility heatmaps. Heatmaps showing the pairwise correlations from Spearman correlation coefficient of peaks compared pairwise. For each gene, peaks are ordered by their importance rank in predicting gene expression and divided into three tiers to determine whether the most important peaks (Tier 1), second-most important peaks (Tier 2), or least important peaks (Tier 3) tend to be most co-accessible. *HLA-DRA*’s Tier 1 peaks have the most co-accessibility of all three tiers. Peaks for *GALC* are moderately accessible across all three tiers, and peaks for *BANK1* tend to be the most co-accessible in Tiers 1 and 2 only.

### GWAS variants are enriched in prioritized peaks

Since only a couple of hundred genome-wide significant genes were considered for each disease, we did not have power to perform an analysis such as stratified LD score regression as a test for enrichment of GWAS signal in important ATAC peaks [46]. Instead, we simply used the aggregated peak ranks to rank the ATAC peaks from each model from most to least important, and determined which peak harbored the lead GWAS variant, shown in Figure 6 (see Methods). This analysis was performed across the three diseases with a total of 478 GWAS gene-to-variant pairs listed in Table 3 evaluated both jointly (Figure 6A) and independently (Figure 6B). There was a strong general trend the GWAS variants to be found in the top 25^th^ percentile of ranks. Although IBD GWAS variants are largely believed to be functional within the gut [47] (either in resident immune or epithelial cells), this result indicates substantial enrichment of regulatory signal in the vicinity of peaks that regulate gene expression also in circulating immune cells. Actual evidence that regulatory peaks correspond to GWAS signals located within accessible chromatin will require experimental methods such as genome editing associated with multiomic profiling.

**Figure 6.**
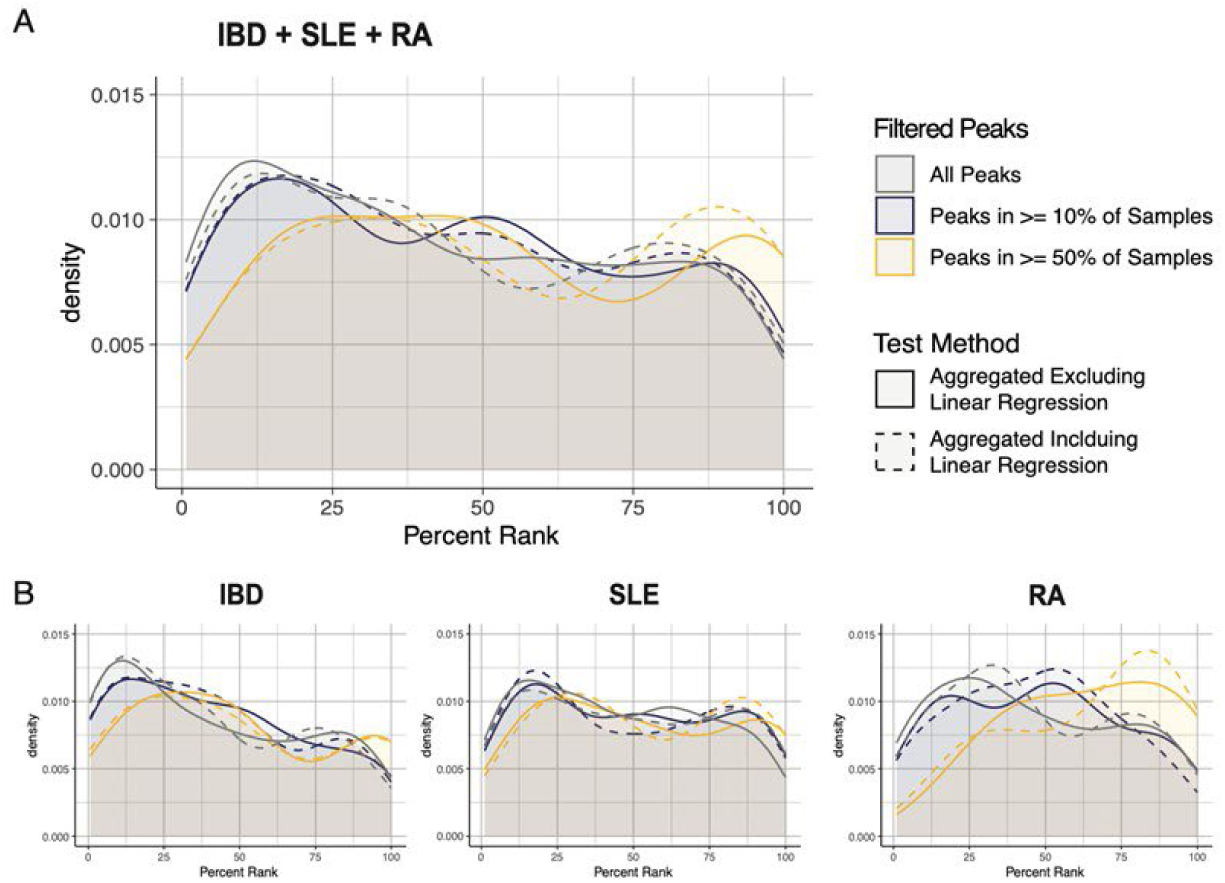
GWAS variants are enriched in important peaks. Density plots illustrate the rank of the ATAC peak used for gene expression prediction which contains the GWAS variant for all traits (A) and stratified by each trait (B).

**Table 3.** Number of GWAS Gene-to-Variant Pairs in Testable ATAC Peaks.

| Trait | All Peaks | Peaks in $\geq 10\%$ Samples | Peaks in $\geq 50\%$ Samples |
| --- | --- | --- | --- |
| IBD | 144 | 114 | 36 |
| SLE | 216 | 162 | 33 |
| RA | 129 | 88 | 19 |
| IBD + SLE + RA | 478 | 354 | 85 |

The enrichment was strongest for the more inclusive peak sets (gray and blue lines in Figure 6), implying that cell-type specific peaks are more likely explaining the GWAS signal than peaks present in at least half the samples (yellow lines). Similar enrichment profiles were observed whether or not linear regression models were included in the aggregation (solid versus dashed lines). Notably, the strongest enrichment was observed for IBD, perhaps reflecting that two thirds of the samples were from donors with this condition. SLE, an autoimmune condition mediated by immune cell dysfunction [48] also shows a peak between the 10^th^ and 25^th^ percentile of importance ranks (Figure 6B), but that was not the case for RA despite the role of immune cells in that class of autoimmunity [49]. It is possible that GWAS variants for RA are more active in the inflamed synovium and associate with peaks specific to that tissue environment [50,51].

## Discussion

Since a large proportion of GWAS variants reside in noncoding regions, understanding their associations with chromatin data has become essential to deconvoluting their impacts on gene regulatory mechanisms [8]. Previous studies and tools predicting gene expression are based on the incorrect assumption that increased chromatin accessibility implies increased gene expression, which has not always been reported to be the case [6,52]. With the advancement of single nuclear RNA and ATAC sequencing from the same cell, we have the unique opportunity to further decipher the influence that the chromatin landscape has on the transcriptome using machine learning so as to identify which peaks are important for explaining gene expression, how well they explain gene expression, and whether GWAS variants tag loci in ATAC peaks tend to be associated with regions important for gene expression variability.

Until recently, investigations of the interplay between chromatin accessibility and gene expression had focused on positive linear correlations. Two recent studies used Poisson regression modeling to assess relationships between chromatin accessibility and gene expression, showing good power to identify regions of chromatin accessibility which are important for explaining gene expression. Both studies implemented models at the single cell level, potentially incorporating noise from drop-outs, and although applied to diverse datasets they compensated for small sample numbers by utilizing the high technical replication in single cells. We chose instead to perform analysis on pseudobulk data ensuring that most of the signal is among individuals and cell types, using a rigorous cross-validation procedure to evaluate model performance. Lack of feature selection of ATAC peaks can also contribute to noisiness, and result in redundancy of correlating peaks being difficult to deconvolute when only one peak may be important. For instance SCARlink utilizes the entire region of chromatin accessibility without peak calling [21]. In contrast SCENT does use MACS2 called peaks, however this study’s implementation binarizes the peaks to be evaluated as open or closed when evaluating the relationship between chromatin accessibility [20]. This removes important biological information of the strength of accessibility, which not only is helpful for considering signal to noise ratios but also for retaining variation across samples and cell types.

Our utilization of machine learning suggests a more nuanced and complex relationship between chromatin accessibility and the regulation of gene expression. We found that linear regression models performed relatively poorly in comparison with three related machine learning techniques. Our analysis employed random forest regression, XGBoost and LightGBM, all of which are sophisticated tree-based machine learning methods known for their proficiency in capturing complex, nonlinear relationships, and potential interaction effects among peaks. Despite their distinct methodologies, these models performed similarly to each other. Because each model has its own strengths and weaknesses, which could lead to performance variances across different scenarios, we elected to aggregate results from all three models. By doing so, we aimed to mitigate the influence of individual model variances and attain a more dependable estimate of predictive accuracy and importance of each peak.

We compared our predictive models against ArchR’s “gene score” predictions, which infers gene expression by summing the total chromatin accessibility around a gene. Our models outperformed ArchR in predicting gene expression more accurately, despite using a subset of ATAC peaks within a gene’s cis-regulatory region. Notably, the cross-validation R^2^ metric provides a more robust and reliable metric than the traditional squared correlation and strongly emphasizes the advantages of the machine learning approach. While ArchR’s “gene score” prediction assumes a direct correlation between increased chromatin accessibility and increased gene expression, our models challenge this assumption by revealing instances where this relationship may not hold true while also incorporating evidence that not all peaks contribute to a gene’s expression. Although ArchR offers the advantage of not requiring training data to infer gene expression from ATAC data, caution is warranted when interpreting gene expression inferred from “gene scores”. Given a scenario where one portion of the dataset is multiome and a second portion is snATAC-seq only, using our predictive models to train on the subset of data with snRNA-seq data would presumably provide more reliable gene expression predictions than ArchR. A secondary use that our modeling approach is the ability to rank peaks by importance for predicting gene expression, providing a strategy to prioritize peaks for further biological interpretations.

Our modeling establishes that only a subset of ATAC peaks within a cis-regulatory region independently contribute to gene expression. We observed that on average, 47% of peaks identified in a gene’s cis-regulatory region were sufficient to explain as much gene expression, if not more, than using all the peaks within the same region. This could be due to a variety of reasons. Firstly, despite using gold-standard peak callers, the inherent noise in ATAC data may lead to the inclusion of peaks that are technical noise. Secondly, some peaks may not contribute to gene expression for biological reasons. For instance, a peak may be proximal to multiple genes but functionally relevant to only one. Alternatively, a peak could be associated with epigenetic processes unrelated to gene expression, such as histone modifications, trans-regulatory effects like chromatin looping, or DNA methylation [23, 34, 53].

An important implication of our findings is that peak prioritization may enhance downstream analyses, such as building gene regulatory networks, which rely on linking genes to peaks, or predicting the potential of a transcription factor to regulate a gene if it binds to a nearby peak [54]. Considering all peaks within a region to be relevant will inevitably generate false connections. While using a subset of peaks explained as much gene expression as using all peaks, unsurprisingly no genes were observed to have 100% of their variance explained from our models. It is for example well appreciated that some peaks may be open for business and only functional in the presence of an activated transcription factor, so differential accessibility is not necessarily a signature of differential activity [55,56]. Nonetheless, we observed that the expression of certain genes is explained by chromatin accessibility better than others, suggesting that some genes may rely more heavily on additional gene regulatory machinery beyond chromatin accessibility. It will be interesting to see as more extensive multiome datasets become available whether general rules describing the architecture of accessible chromatin influence on gene expression can be derived.

We further investigated the potential contribution of each peak on gene expression by computing Shapley values, which shed light onto whether a peak may contribute to an increase or decrease in gene expression prediction values. Surprisingly, we found that approximately 50% of the peaks we evaluated contributed to a decrease in gene expression prediction values, negating the widely held assumption that increased chromatin accessibility generally leads to increased gene expression. Notably, univariate linear correlations between each peak and the real gene expression were also assessed, only suggesting 15% of the evaluated peaks to have a negative correlation with gene expression. Shapley values account for interaction effects between peaks, nonlinear relationships, and redundancy, so a large proportion of negative values can be taken as evidence that co-accessibility contributes to correlations between single peaks and nearby transcript abundance. On the other hand, there is some tendency for Shapley values within a cell type to have the same sign, and since the models are assessed across all cell types it is possible that there is an overall tendency for the Shapley values to balance positive and negative signs. In addition, 27% (n = 5,533) of the peaks we assessed were observed to be relevant to more than one gene. Of these peaks, 43% contributed to gene expression in the same direction for all relevant genes (either all positive or all negative contributions), while the remaining 57% simultaneously contributed to an increased predicted gene expression value for one gene and a decreased predicted gene expression value for another. These findings further highlight the complexity of the chromatin accessibility landscape, and the variable effects it has on gene expression.

Discerning the functional relevance from prioritizing peaks additionally provides an avenue for inferring genetic effects on gene regulation. We identified which peaks tended to harbor a GWAS tagged variant and found that the top 25 percentile of peaks ranked to be important for explaining gene expression were enriched with GWAS variants, specifically for IBD and SLE. This is despite the expectation that many of the lead SNPs are proxies for the true causal variant in high linkage disequilibrium, yet is consistent with the widespread belief that GWAS tagged loci play a mechanistic role in gene regulation and hence act as eQTL [57–59]. That this is often not the case was made clear by Mostafavi et al. (2023)[60] who showed that eQTL and GWAS variants have different statistical properties and quite often do not colocalize. Consequently, the modest degree of enrichments observed in Figure 6 were not unexpected. Although this dataset is relatively small, an enticing application of snATAC-Express on a larger dataset would be to evaluate the enrichment of GWAS variants within prioritized ATAC peaks among individuals within each cell type.

Limitations of this study include the size of this dataset, which contains just 29 individuals and 18 distinct cell type populations. Datasets consisting of several hundred individuals would enable more refined calculation of ATACseq peak weights for specific cell types: our current models are largely driven by chromatin accessibility differences among cell types. Larger samples will also presumably be less susceptible to biases driven by outlier samples, though the cross-validation approach should limit dependency on such samples. This dataset is unique, in that it contains samples from a cohort of individuals with Crohn’s disease and healthy individuals, all with African American ancestry [61]. Given the current scarcity of multiome datasets which are similar, assessment of generalizability of our predictive models was limited to stratified k-fold cross validation. However, as multiome data becomes more widely available and datasets become larger, these models can be evaluated on additional publicly available data. Once sufficiently large multiome datasets are available to train models unique to each cell type, we would expect that fewer peaks will be found to be critical for each cell type and that enrichment for GWAS signals in the most important peaks may increase. Another limitation of predicting gene expression using snATAC-seq data with our models is that it requires paired snRNA-seq data for training and testing, making it not feasible to be used on isolated snATAC-seq data without another dataset of similar characteristics that has snRNA-seq data associated with it. Multiome assays are expensive and time consuming, so this method could provide a solution to perform multiome on a subset of a cohort and only scATAC-seq on the remaining cohort, using the multiome cohort for training to predict the gene expression on scATAC-seq only samples.

The results presented here describing the chromatin landscape and prioritization of ATAC peaks which are important for explaining gene expression also require experimental validation. For instance, eCROP-seq or Perturb-seq experiments of perturbing variants or eQTL located in ATAC peaks found to be important for explaining gene expression could provide further evidence as to whether those features play a mechanistic role in the target gene’s expression, or, whether another gene’s expression is affected instead [62, 63]. Additional functional annotations such as from ChIP-seq experiments to determine whether a transcription factor or other DNA binding protein binds to ATAC peaks of interest [64], or chromatin conformation capture experiments like Hi-C could also elucidate gene regulatory mechanisms and machinery as they relate to prioritized ATAC peaks [65].

## Conclusions

Using snATAC-Express, a machine learning pipeline leveraging multiome data to infer transcript abundance from chromatin accessibility, we demonstrate that approximately half of the ATAC peaks in the vicinity of a gene contribute to models of gene expression. These models, trained on pseudobulked snRNA-seq data and validated with the cross-correlation coefficient of determination, significantly outperform gene scores that simply aggregate open chromatin. We find that the peaks prioritized for making the greatest contribution to explaining gene expression tend to exhibit increased co-accessibility with one another, and they are enriched in GWAS variants for at least one inflammatory and one autoimmune disease. Our study emphasizes the feasibility and utility of predicting gene expression using a subset of ATAC peaks using machine learning models which are flexible to interaction effects and nonlinear relationships.

## Methods

### Single nuclear RNA-seq and ATAC-seq dataset

The single nuclear RNA-seq and ATAC-seq dataset is from peripheral blood mononuclear cells (PBMC) and consists of 29 individuals and 18 identified cell types [60]. Libraries were made using the Dual Index Kit TT set A (part # 1000215) and sequenced at the Georgia Tech Molecular Evolution Core in accordance with 10X Genomics’ recommendations. The fastq files were aligned using cellranger-arcv2.0 software, with the refdata-cellranger-arc-GRCh38-2020-A-2.0.0 reference [66]. Library generation, data processing, quality control, and cell type identification steps are described in Brown *et al*. (2024)[61]. Single nuclear data was pseudobulked by donor and cell type, resulting in a total of 342 pseudobulked samples after exclusion of samples with fewer than 25 cells. For snRNA-seq data, raw count data was first converted to counts per million (CPM) after summing the counts for all cells of a cell population for each donor and dividing by the total number of reads per donor. This value was then converted to CPM by multiplying by a scale factor of 1,000,000, and then log2 transformed by computing log2(CPM + 1). For snATAC-seq data, the peak counts identified from MACS2 were summed for all cells of a cell population and divided by the total number of peak counts per donor [67]. This value was then converted to CPM by multiplying by a scale factor of 1,000,000, and then log2 transformed by computing log2(CPM + 1). Pseudobulked snRNA-seq and snATAC-seq values were used for predictive modeling.

### Gene and ATAC peak selection

A total of 502 genes were selected for predictive modeling. Genes tagged by GWAS variants reported to be genome-wide significant for IBD, RA, and SLE were obtained from the GWAS Catalog (https://www.ebi.ac.uk/gwas/; accessed January 2024) [68]. For modeling, three criteria were required to be met by the gene: 1.) it must have a gene expression value across all pseudobulks in our dataset greater than zero, 2.) it must have at least one ATAC peak in one sample within its cis-regulatory region that contains the tagged GWAS variant, and 3.) it must have at least three ATAC peaks within its cis-regulatory region. Cis-regulatory regions for each gene were defined as 100kb upstream and downstream of the gene’s transcription start site and end site, respectively, also including the introns and exons of the gene body [22]. Gene coordinates were obtained from Gencode Release 41 (GRCh38.p13) [69]. ATAC peaks observed to overlap this region were assigned to the gene for model training.

### Predictive modeling

The code for the following methods and generation of figures can be found on our GitHub repository: https://github.com/GibsonLab-GT/snATAC-Express including a Tutorial: https://github.com/GibsonLab-GT/snATAC-Express/blob/main/tutorial.ipynb. The expression of each gene was trained on ATAC peaks within the cis-regulatory region for that gene, using pseudobulked values. Four types of models were implemented: linear regression, random forest regression, XGBoost, and LightGBM. Linear regression and random forest regression modeling was implemented using functions from scikit-learn [70]. XGBoost was implemented using the xgboost python package, and LightGBM was implemented using the lightgbm python package [71,72]. Two rounds of predictive modeling were implemented, as shown in Figure 1C.

First, models were trained on pseudobulked snATAC-seq peaks three ways: all peaks within the gene’s cis-regulatory region, peaks in at least 10% of samples, and peaks in at least 50% of samples. For each of these three ways, two to three methods of feature importance ranking assessments were implemented: model weights (machine learning methods only), permutation ranking, and drop column ranking, for a total of 11 models. K-fold cross validation was implemented for each implementation, and the data was divided into 3 folds, with 5 splits per fold, resulting in a total of 15 splits across all folds. The R^2^ values for each k-fold cross validation step per model implementation were aggregated by computing the average cross validation R^2^ value, which is reported in the violin plots for Figure 1D and Supplementary Figure 1A-B. In addition, the peak ranks for each k-fold cross validation implementation were aggregated by averaging the weighted values. The top 95% cumulatively important peaks were identified by averaging the peak ranks across all implementations from the initial modeling implementation (after a Z-score conversion), and then the peaks whose cumulatively summed weights (from highest to lowest) explained approximately 95% of the total weights for all peaks were selected for second round modeling, stratified only by whether the implementation was from including all peaks, peaks in at least 10% of samples, and peaks in at least 50% of samples. The second round of modeling underwent k-fold cross validation in the same way, and the R^2^ values and peak ranks were aggregated the same as the initial round of modeling. Hyperparameter tuning was performed prior to k-fold cross validation using the gridsearch() function from scikit-learn for the three machine learning models [70]. No meaningful bias was introduced by including the second step; all of our results hold for models without it but should be slightly more accurate.

### Peak ranking and GWAS enrichment

For each model, peaks were ranked from most to least important in explaining gene expression. Because some ranking methods included negative numbers (such as drop column ranking) while others did not, a z-score was computed for each method’s peak ranks, which were averaged across methods for aggregation. Peak ranks were normalized to be between 1 and 100 by computing the peak rank divided by the total number of peaks multiplied by 100, where 1 indicates the most important peak, and 100 indicates the least important peak. Peaks which overlapped the SNP for the GWAS variant to gene pair were determined by whether the SNP’s gene coordinated overlapped the ATAC peak coordinate. Note that this approach assumes that the reported GWAS variant is the causal variant, which is often not the case due to high LD, and consequently there is no expectation that all GWAS peaks will lie within the most important ATAC peaks. Supplementary Figure 5 shows which peaks tend to contain the GWAS variant across all models, for models using all peaks, peaks in at least 10% of samples, and peaks in at least 50% of samples.

### Pathway enrichment

The 500 genes selected from GWAS tagged variants for IBD, RA, and SLE which also passed filters for generating models using peaks in at least 10% of samples were used for pathway enrichment analysis. Pathways enriched in these genes were identified using ToppFun, which is part of the ToppGene Suite [73]. Queries were performed against multiple databases, including BioCarta Pathways, Canonical Pathways, KEGG, Pathway Interaction Database, Reactome, and WikiPathways. A Bonferroni p-value correction was applied for multiple comparison control using a 0.05 significance threshold.

### Inferred ATAC peak impact on predicted gene expression

To infer the contribution of the ATAC peaks on gene expression prediction in the trained models, we computed Shapley values using the shap Python module [37–39]. During K-fold cross validation, Shapley values were determined within each fold for each sample and peak, and then aggregated by computing the mean Shapley values across folds. For each peak, we compared it’s Shapley values against the accessibility values to determine whether a positive or negative correlation was observed by assessing the slope for the line of best fit using sklearn’s LinearRegression() function as shown in Figure 4C [70]. In addition, we compared each ATAC peak against the gene it is being used to predict and also determined whether a positive or negative correlation was observed by assessing the slope for the line of best fit using sklearn’s LinearRegression() function. The Shapley value approach accounts for interaction effects between peaks, nonlinear relationships, and redundancy in the model context. A negative Shapley value indicates that increased accessibility at that peak, in the context of other peaks in the model, contributes to decreased predicted gene expression This could reflect: (i) binding sites for transcriptional repressors, (ii) competitive binding between activating and repressing factors, or (iii) complex regulatory logic where peak combinations determine outcomes.

### Co-accessibility analysis

The Spearman correlation was computed using the Python Pandas package for peaks in all genes across two categories: the first including all peaks, and the second only including the top 95% cumulatively important peaks that appear in at least 10% of the pseudobulked samples [74]. The *n* × *n* correlation matrices were subdivided into three equal parts to analyze specific triangular regions along the diagonal. This was accomplished by generating three masks, where each mask targeted a square section by first calculating a side length 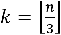 and then distributing the squares along the matrix’s diagonal. Each mask covered a different triangular region: top left, middle, and bottom right, ensuring an equal partition of the matrix. An additional step involved applying an upper triangular mask to each, removing duplicate values from the analysis. In the case in which *n* isn’t divisible by three, the remaining peaks are split between the middle and bottom right square.

For matrices in the 95% cumulatively important peaks that appear in at least 10% of the pseudo-bulked samples category, the values in each region were scaled between 0 and 1 using minmax scaling, and the average of the values in each of the regions was computed. This average was then classified into three bins—low, medium, or high—corresponding to three bands: 0 to 0.33, 0.34 to 0.66, and 0.67 to 1. The permutations of these three bins across the three masks were finally counted across all genes to assess an underlying structure of the data.

### Comparison with ArchR gene scores

Gene score values were obtained from ArchR using ArchR’s addGeneScoreMatrix() function [17]. Like the true gene expression previously, these values were pseudobulked the same way to obtain log2(CPM +1) counts. For each gene, these values were compared with the real gene expression values. The linear least squares R^2^ values were obtained for each gene (the same was done for the machine learning predictions aggregated across k-fold cross validations versus the real gene expression values), in addition to the 1-(RSS/TSS) R^2^ values. To ensure appropriate comparisons for ArchR, log2(CPM +1) counts of the gene score values were compared to the true gene expression log2(CPM +1) counts, and the z-scores of these values were also compared. Heatmaps were generated for comparison using the z-scores of log2(CPM +1) counts for all three versions, and Ward’s clustering was employed on the genes and samples for the real gene expression values only. The order of genes and samples from clustering using the real gene expression values were preserved for generating heatmaps for predicted values from machine learning models (trained on peaks in at least 10% of samples) as well as ArchR’s gene scores.

## Declarations

### Ethics Approval

All biopsies were obtained from patients with inflammatory bowel disease or undergoing endoscopy for other reasons under a protocol approved by the Emory University Institutional Review Board (IRB) with further permission for genomic analyses provided by the Georgia Tech IRB (Protocol H11286). Written informed consent was obtained from all subjects and/or their legal guardian(s). All sampling and analyses were performed according to the relevant guidelines and regulations pursuant to the Declaration of Helsinki.

### Data Availability

All snATAC-seq and snRNA-seq data are available through the Gene Expression Omnibus through accession GSE244831. All code and supplementary supporting data is provided at Maggie Brown’s Github repository, https://github.com/maggiebr0wn/scMultiome-Crohns-Disease.

### Author Contributions

MB performed or supervised all statistical and bioinformatics analyses, designed the snATAC-Express pipeline, assembled the figures, and wrote the draft manuscript. AF performed the analyses reported in Figure 5. AD provided clinical study management and with FS and VK processed the samples for multiome analysis. SK oversaw patient care and sample acquisition and was principal investigator for the project. RDW provided statistical advice and interpretation. GG conceived the study, helped write the manuscript, and supervised MG and AF.

## Acknowledgements

We thank all donors for their consent to participate, particularly during the COVID-19 pandemic, and members of the Gibson and Kugathasan labs for discussions.

---

**Supplementary Figure 1.**
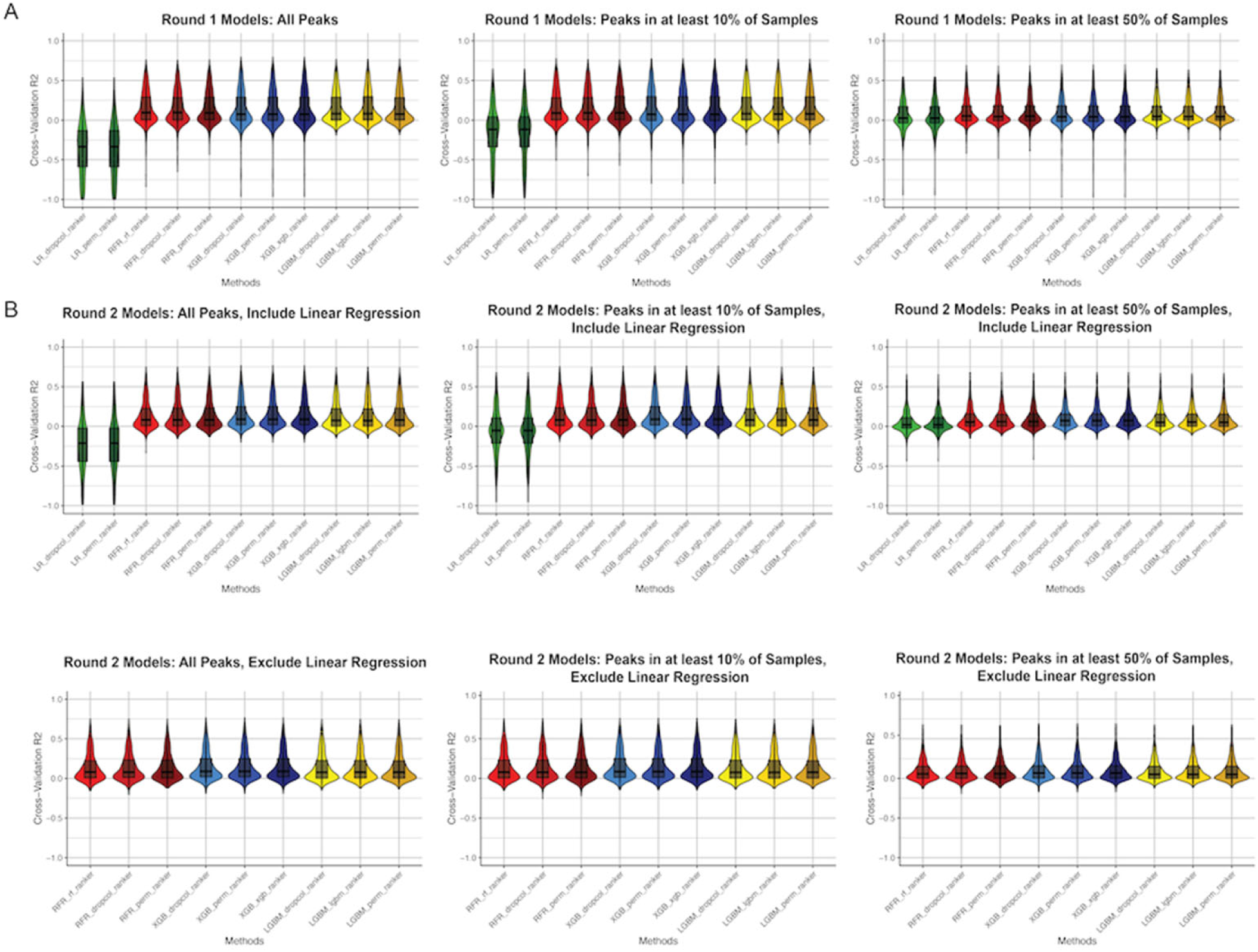
Evaluations of all models. (A) R2 values aggregated from k-fold cross validation across all Round 1 models, computed as 1-(RSS/TSS). (B) R2 values aggregated from k-fold cross validation across all Round 2 models, computed as 1-(RSS/TSS) including linear regression (top) and excluding linear regression (bottom). RSS, residual sum of squares; TSS, total sum of squares.

**Supplementary Figure 2.**
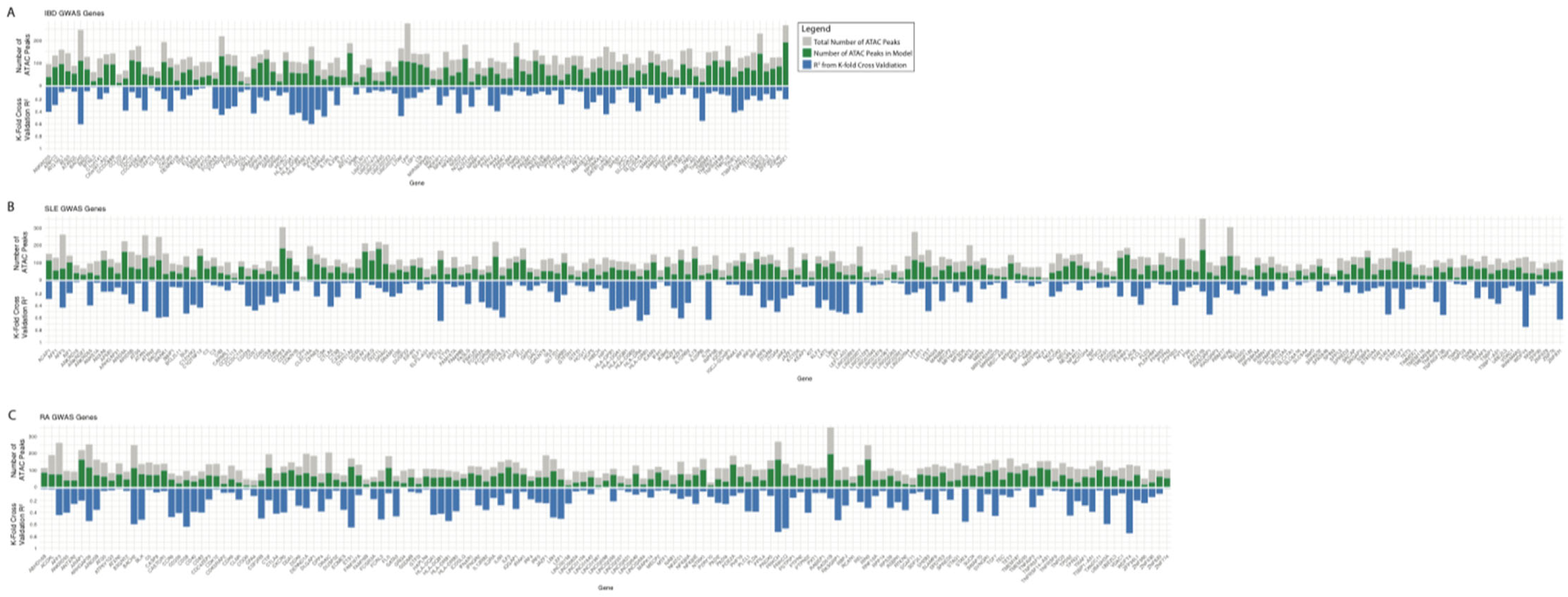
Atlanta plots for all GWAS traits. Atlanta plots of GWAS tagged genes from IBD (A), SLE (B) and RA (C). The gray bars represent the total number of peaks within the gene’s cis-regulatory region, and the green bars represent the number of peaks in the region used to trained on for gene expression prediction. The blue bars represent the amount of gene expression explained, as the ensembled R2 value from k-fold cross validation. Sets of genes are clustered by pathway and arranged from left to right alphabetically.

**Supplementary Figure 3.**
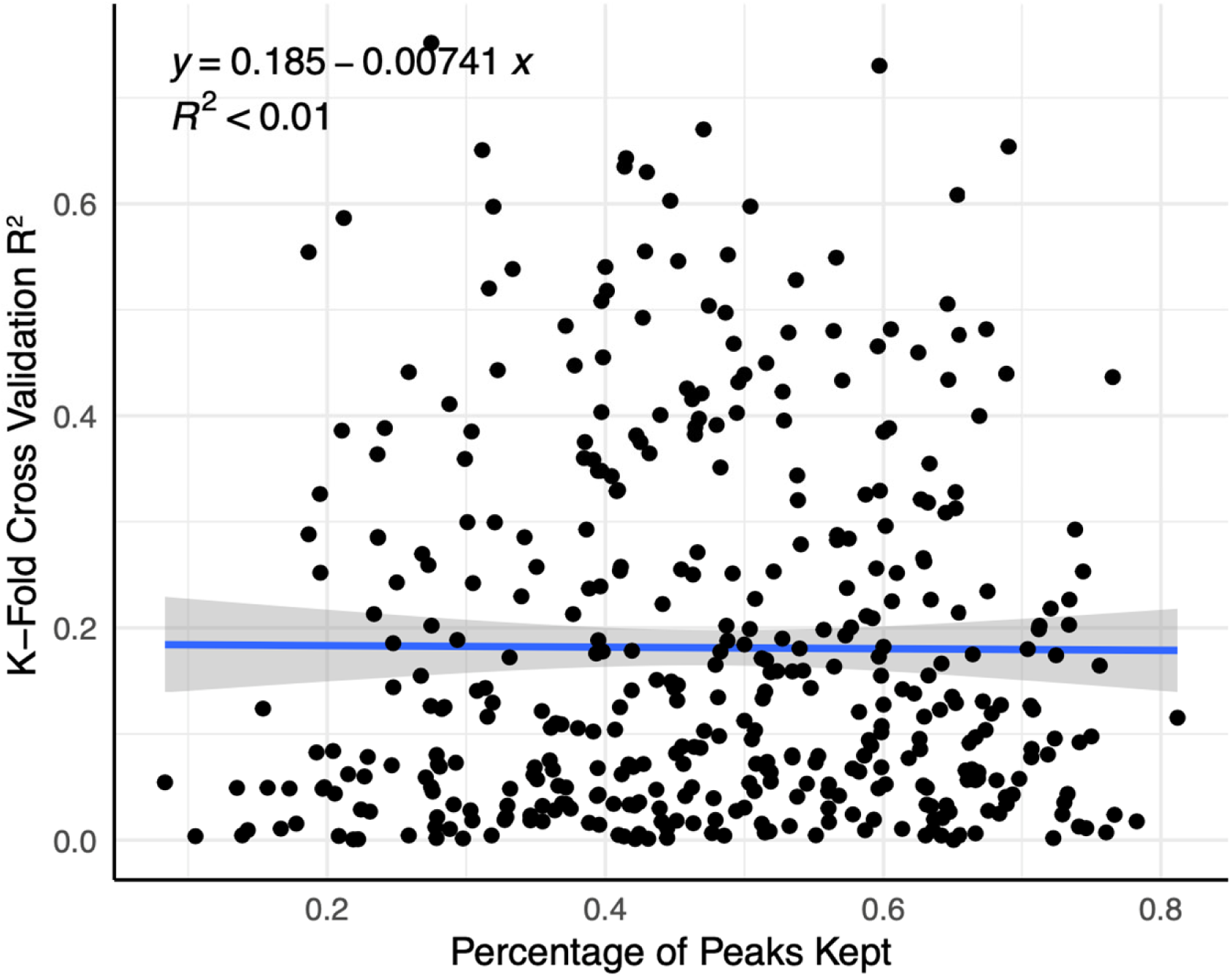
Comparison of R2 values versus percentage of retained peaks. Each gene’s aggregated k-fold cross validation R2 value from machine learning models trained on the top 95% cumulatively important peaks present in at least 10% of samples, versus the percentage of the peaks present in the gene’s cis-regulatory region used for training.

**Supplementary Figure 4.**
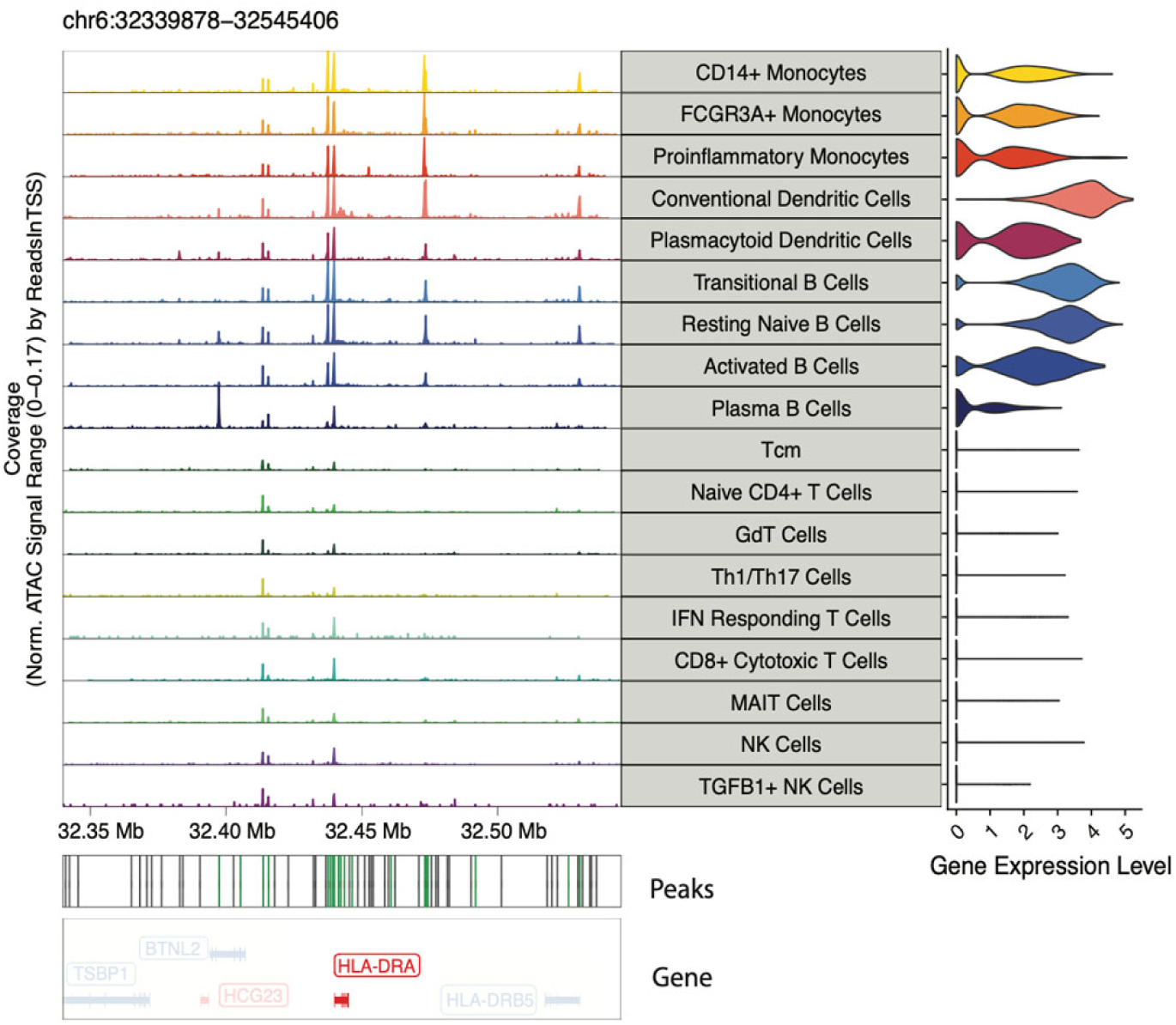
A subset of HLA-DRA peaks explain gene expression. A track plot for gene HLA-DRA, which shows 61 ATAC peaks within the region across circulating immune cell populations, of which 19 peaks (shown in green) are used for gene expression prediction.

**Supplementary Figure 5.**
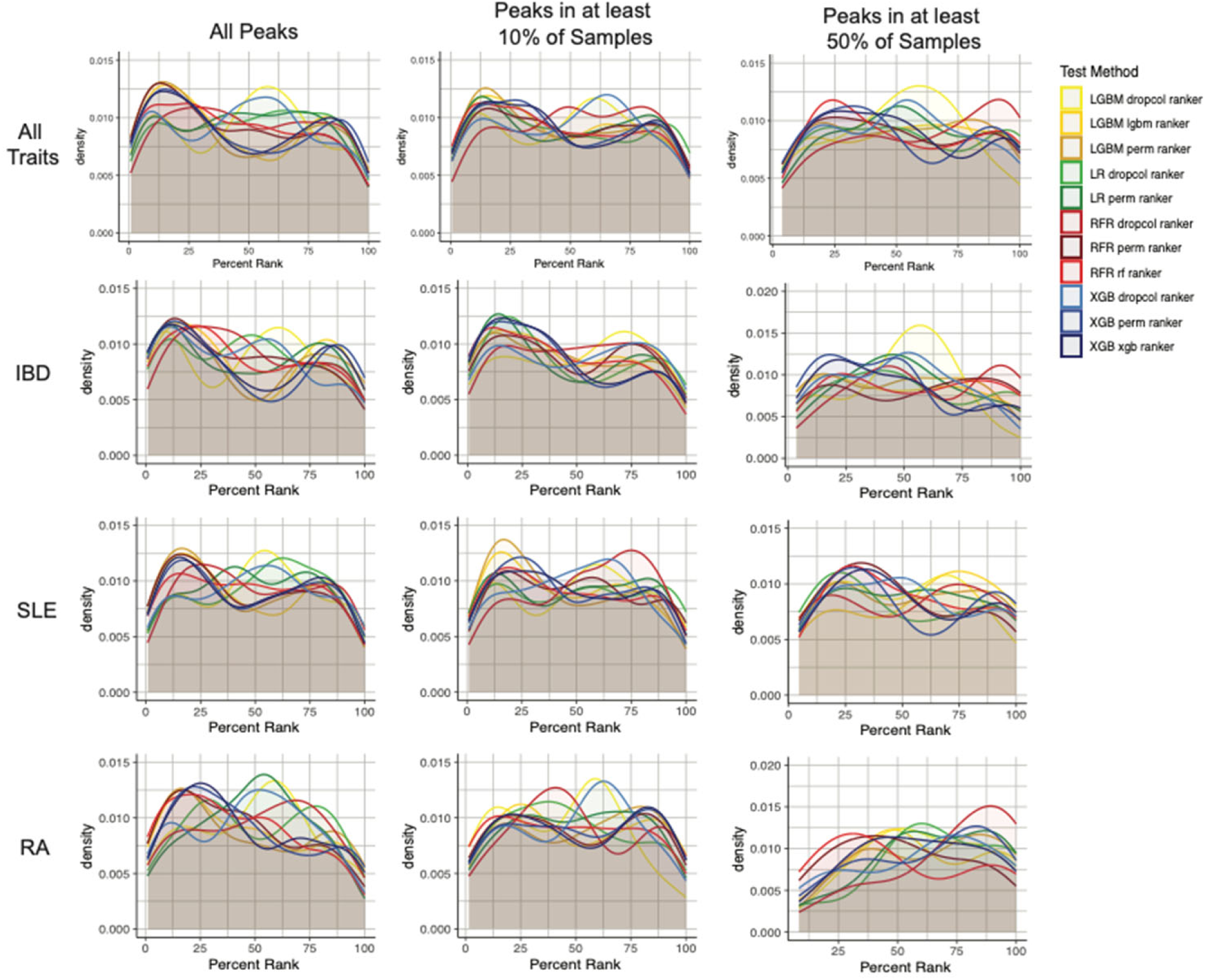
Enrichment of GWAS signals near priority ATAC peaks. Density plots depict the rank of the ATAC peak used for gene expression prediction which contains the GWAS variant for all traits on interest, stratified by pipeline version and modeling method.

## List of Supplementary Tables (Available upon Request)

<u>Supplementary Table S1.</u> R-Squared Statistics of Model Fits

Values represent the mean and median R2 computed as the linear regression fit or coefficient of determination, for models with or without peaks identified by linear regression, at three levels of inclusion, and compared with ArchR fits.

<u>Supplementary Table S2.</u> Gene-to-Peak Correlations with Shapley Values

The table lists the number of instances of positive or negative correlation between observed gene expression and Shapley values from aggregated models, for SLE, IBD and RA associated genes.

<u>Supplementary Table S3.</u> Number of Genes per ATAC Peak

Tabulation of the number of instances where an ATAC peak was associated with expression of from 1 to 8 or more genes.

<u>Supplementary Table S4.</u> Patterns of Co-Accessibility for the Three Diseases

Tabulation of number of instances of each of a dozen peak co-accessibility patterns for SLE, IBD, and RA associated genes.

